# Cloning and *In silico* characterization of Heat shock factor (Hsf2) from Wheat (*Triticum aestivum L.*)

**DOI:** 10.1101/2020.04.17.046094

**Authors:** Kavita Dubey, Suneha Goswami, Narendra Kumar, Ranjeet R. Kumar, Ravi RK Niraj

**Affiliations:** Division of Biochemistry, ICAR-Indian Agricultural Research Institute, New Delhi-110012; IMS Engineering College,Ghaziabad,Uttar Pradesh −201009; Amity Institute of Biotechnology, Amity University Rajasthan, Jaipur

**Keywords:** Wheat, heat shock factors HSF), *in silico*, Thermal stress

## Abstract

A gene that encodes for Heat Shock Factor (Hsf) was characterized and designated as TaHsf2 (Gen Bank ID: KP063542) from wheat. The genomic sequence was found to be 1551bp long with open reading frame of length1123bp. The amino acid sequence so concluded showed to have HSF domain having high degree of similarity (homology) with other HSF coding gene of related species. We have reported in-silico three-dimensional model of TaHsf2 and evaluation of the predicted model, analysed with various tools. The sudden climatic variations over the years especially elevation in temperature has adversely affected the growth and productivity of winter wheat (*Triticum aestivum* L.). A very less number of HSFs is being reported and characterized in wheat and the data so available for wheat calls for investigation and characterization of more HSP’s coding HSFs. In our present study we have cloned one of the novel HSFs and have characterized it through *in silico* tools.

## INTRODUCTION

Heat stress is one of the major abiotic stresses which adversely affects the growth and productivity of crops all around the world. The climatic variations put the crops under two categories of stress: biotic and abiotic stress. Both biotic and abiotic stresses equally contribute to the changes in the biological and physiological states of the plant and have adverse consequences. Among the abiotic stress thermal stress, especially the elevation in temperature during the pollination and grain-filling stages has adverse effect on the growth and yield of wheat (*Triticum aestivum* L.) Wheat (*Triticum aestivum* L.) tolerate heat stress to varying degrees at different phenological stages, but heat stress during the reproductive phase is more harmful compared to vegetative phase due to the direct effect on grain number and dry weight (Wollenweber *et al*., 2003).All plants adapt up with the changes in encompassing temperature push by bringing changes in membrane fluidity, perturbation in metabolism, protein affirmation and assembly of the cytoskeleton(Ruelland and Zachowski, 2010). The cycle of responses actuates versatile forms counting expression of heat shock proteins, until modern cellular balance comes. The temperature above the optimum causes irreversible damage or injury resulting in ‘heat stress’ which is deleterious for the growth and development of crops (Wahid *et al*., 2007).

The duration and level of heat stress adversely affects the protein metabolism, inhibits protein accumulation, and induction of certain protein synthesis (P. Monjardino *et al*. 2005; Y. He and B. Huang; 2007).The plants brings changes at molecular as well as biochemical level to adapt with the elevation in temperature. At molecular level under heat stress there is transcription and translation of small set of proteins,called heat shock proteins(HSPs) (E. Vierling 1991; C. Blumenthal et al. 1990; M. Krishnan et al.1989).There are many reports which proves that expression of HSPs are mainly regulated by HSFs under heat (Scharf et al., 2012).These transcription factors are considered to be the terminal component of signal transduction pathway that are responsible for expression of genes involved in abiotic stresses (Nover et al., 2001).The HSFs maintain the homeostasis of protein folding and brings the tolerance level under thermal stress.

The characterization of HSFs will provide the basis for studying their role in abiotic stress at different levels. In case of wheat some of the HSFs is being reported and characterized, thus we have investigated one of the HSFs from wheat and have characterized it at *in silico* level.

## MATERIAL AND METHODS

### Plant material

The seeds of thermo-tolerant wheat cultivar HD2967 were sown under controlled conditions of (22°C, 16h/ light/8h dark cycle) in National Phytotron Facility of IARI. Seedlings sown developed after fourteen days were subjected to thermal stress at 42°C for 2 hour and sample for nucleic acid extraction was collected in liquid nitrogen and frwas collected in liquid Nitrogen and freeze at −80°C until nucleic acid extraction.

### Isolation and Cloning of *Hsf* in wheat

The whole RNA was separated from 100 mg leaf tissue utilizing TriZol reagent (Invitrogen, Carlsbad, CA, USA) taking after the manufacturer’s enlightening. The surrender and quality of DNAase (Promega Life Sc, India) treated RNA were decided by Nanodrop 1000 (M/S Thermo-Scientific Enterprise, USA) and 2% agarose denaturing gel electrophoresis in MOPS {3-(N-morpho-lino) propane sulfonic corrosive)} buffer, individually. The cDNA was synthesized utilizing 1 µg of RNA with 200 U l−1 invert transcriptase Superscript III (Thermoscientific), 10 mM dNTPs, and 250 ngoligo (dT). For planning the forward and switch groundworks of CA quality, nucleotide grouping of distinctive vegetable species of angiosperm CAs were downloaded and adjusted. Appropriately two deteriorate preliminaries, i.e. D-FP and D-RP were utilized (Table 1). The intensification item was gel decontaminated with Gel Extraction Unit (Qiagen) and cloned into pGEM-T simple vector (Promega, USA) in *Escherichia coli* strain M13 for sequencing.

### Sequence similarity search and phylogenetic analysis of TaHsf

The homology search of TaHsf sequence was performed using BLASTn (https://blast.ncbi.nlm.nih.gov/Blast.cgi?PAGE_TYPE=BlastSearch). Furthermore, a phylogenetic tree was constructed based on the nucleotide sequence using NCBI tool. **Structural and functional analysis of TaHsf** The 3D structure was predicted using SWISS-modeller using automated mode (Arnold et al., 2006). The modelled structure was subjected to validation and assessment using protein structure and model assessment tools at Swiss-Model server using different estimation patterns. The selected model was validated for packaging quality using Anolea plot (Melo and Feytmans, 1998) while Qmean (Benkert et al., 2009; while PROCHECK (Ramachandran plot) was used to analyze the stereochemical and overall quality of the structure (Laskowski et al., 1993) CDD tool (Conserved Domain Database) from NCBI (http://www.ncbi.nlm.nih.gov/cdd) was implemented for domain and family analysis of amino acid sequence. The ExpasyProtparam tool (http://web.expasy.org/protparam/) was applied for computing the physiochemical characterization of Hsf. The protein-protein inte ++raction study was performed using STRING database.

## RESULTS AND DISCUSSION

### Sequence identification of cloned Hsf

The amplicon measure so gotten after RT-PCR has size of 1.5kb and cloning in pGEMT followed by *Eco*RI restriction (Fig.1) confirms the insertion of our target gene in maintenance vector. The cloned gene was submitted in NCBI database with accession no.KP063542.

**Figure1.**
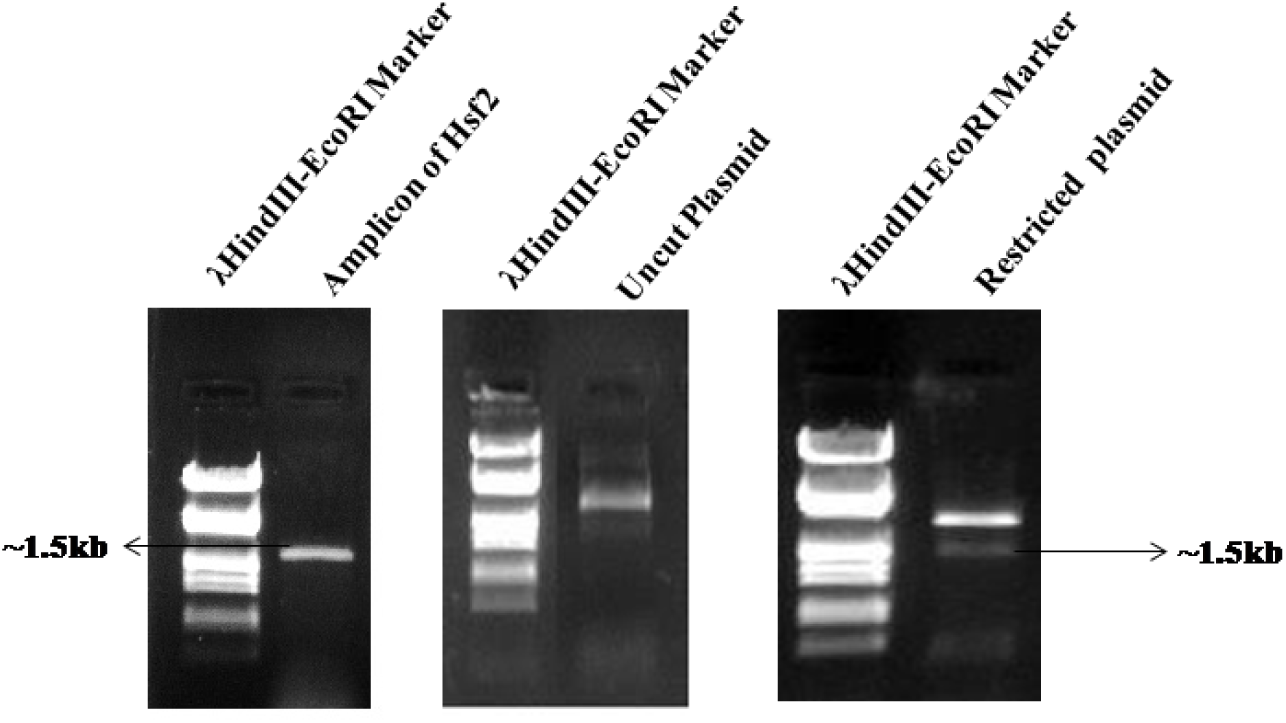
RT-PCR(Reverse transciptase polymerase chain reaction) amplification of HSF2 gene from HD2967 cv. of Triticum aestivum giving amplicon of ~1.5kb with gene specific primers,cloned in pGEMTeasy vector and restriction of plaasmid with EcoRI shows the release of 1.5kb insert checked on 1% agarose gel

### Evolutionary relationship of cloned Hsf with other plant Hsfs

To know the close relatedness of cloned Hsf with other characterized plant Hsf proteins phylogenetic analysis was performed. Phylogenetic analysis revealed that TaHsf has the highest sequence identity and similarity with heat sock transcription factors of *Aegilops tauschii* and unidentified protein of Wheat. (Fig.2).

**Figure2:**
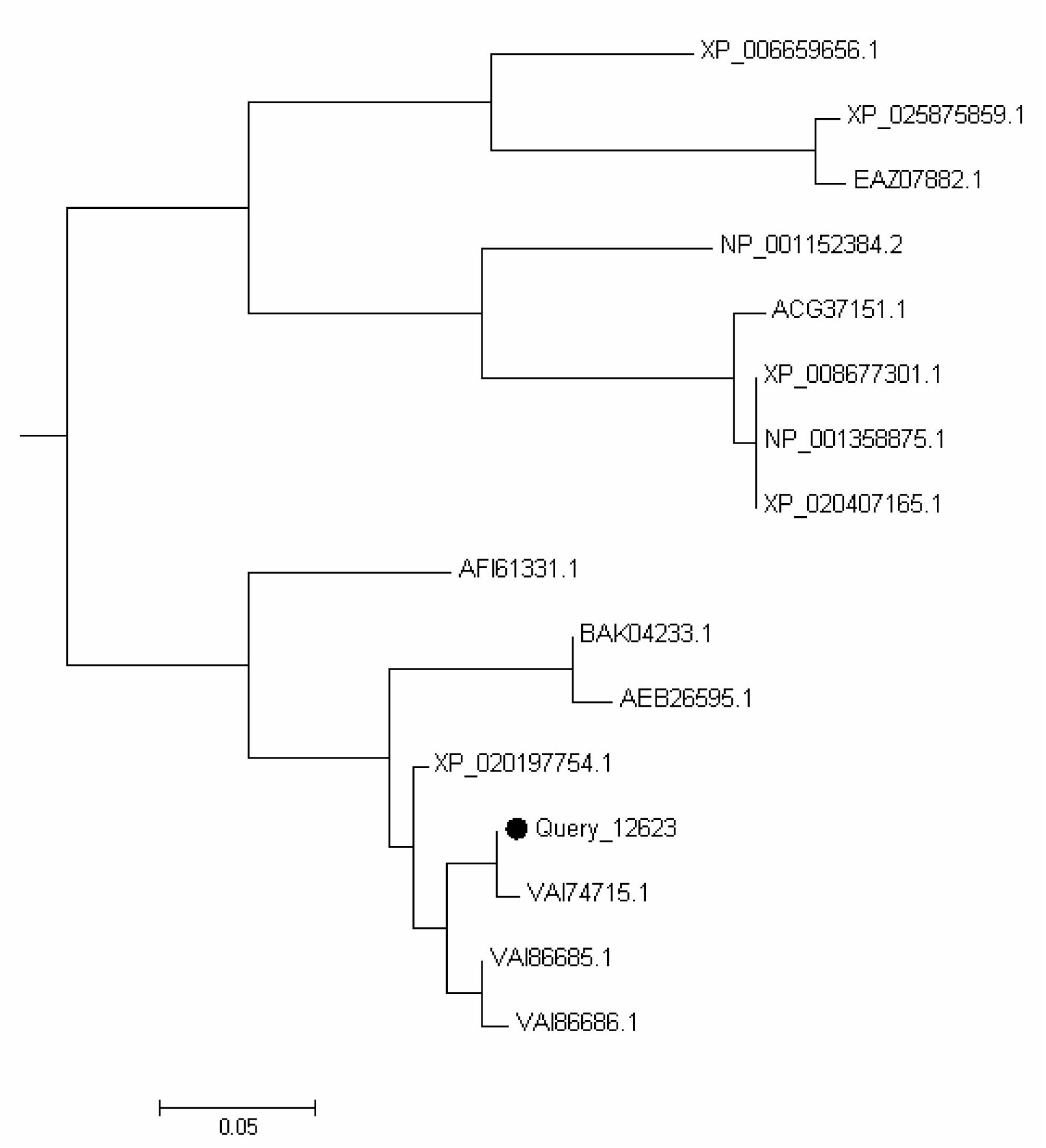
Distance tree for TaHsf2 (Gen Bank ID: KP063542) from wheat using NCBI tool.

### Homology modelling and model evaluation

The structure was modelled using Swiss model online server using automated mode. The structures generated were assessed using Protein Structure and Assessment Tools at Swiss Model server. The selected model (Fig. 3) had the predicted (Zhou and Zhou, 2002) GMQE and QMEAN6 score 0.17 and −0.63 respectively. The conserved domain of was predicted using CD search tool of NCBI that ravel presence of HSF domain (Figure4). PROCHECK (Ramachandran Plot) depicted 86.7% in favourable regions (Fig. 5) and was above the optimal score. The physio-chemical parameter predicted by ExPASy ProtParam tool provide significant information about the composition as well as nature of the protein. Number of amino acids: 374, Molecular weight: 40449.01.The higher value of extinction coefficient (Ext. coefficient 35785) indicates that most of the protein part is occupied with aromatic amino acids, but the value of Instability Index (52.53) more than 40 signifies the instability of protein. The high aliphatic index (Aliphatic index: 65.29) indicates that the protein is thermo stable. GRAVY which is the estimate value for the solubility of the protein is negative indicating that protein is hydrophilic and thus soluble (GRAVY): −0.552). Protein-Protein Interaction of TaHsf network generated by the STRING database based on spring model are shown in Figure6. PPI enrichment p-value is 0.718.

**Figure 3:**
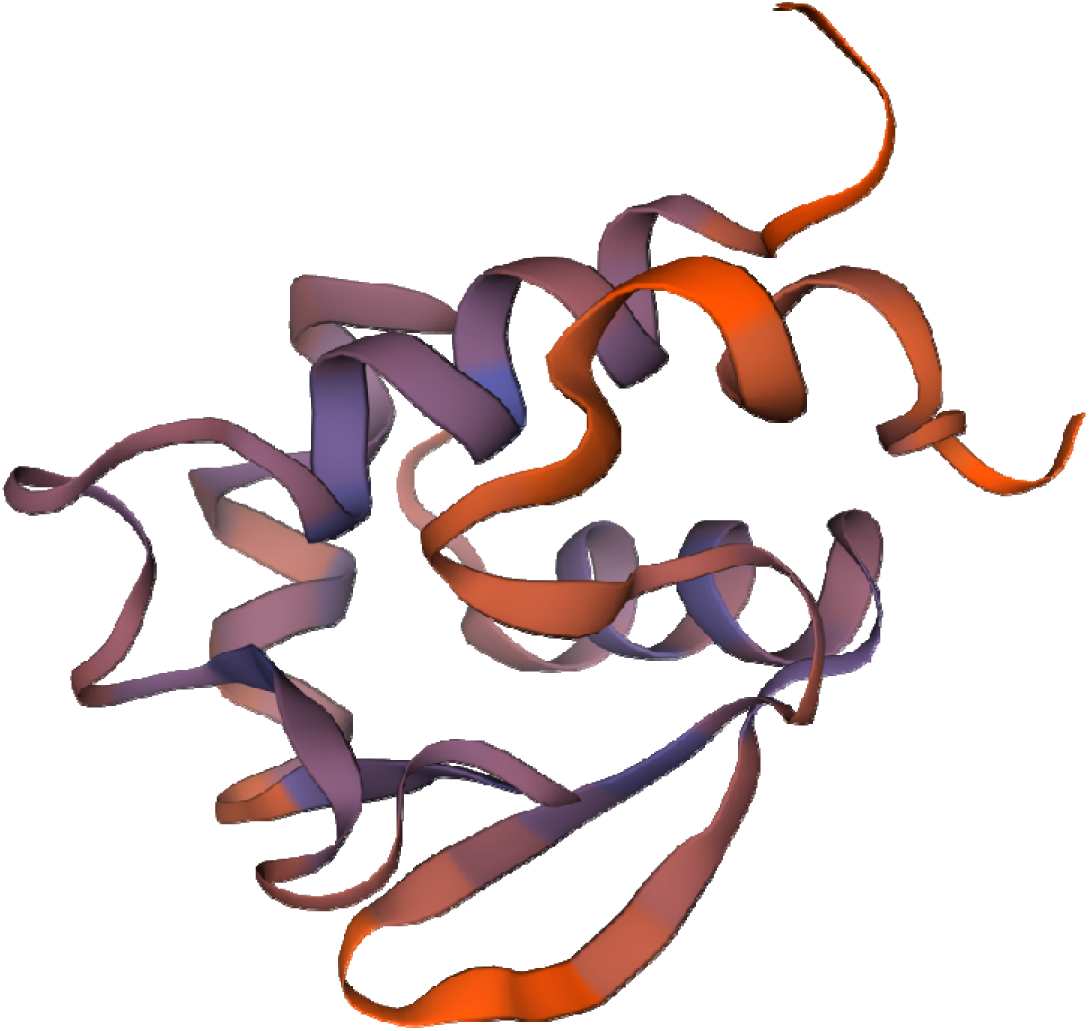
3D modelled structure of TaHsf2 (Gen Bank ID: KP063542) from wheat using NCBI tool.

**Figure4:**
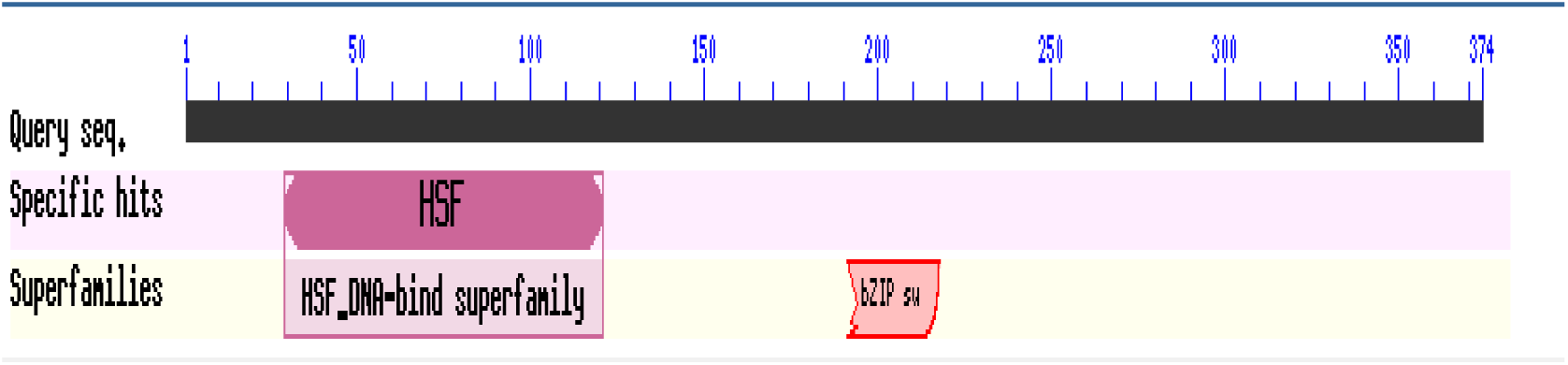
Conserved domain prediction of TaHsf2 (Gen Bank ID: KP063542) from wheat using NCBI CD search tool.

**Figure 5:**
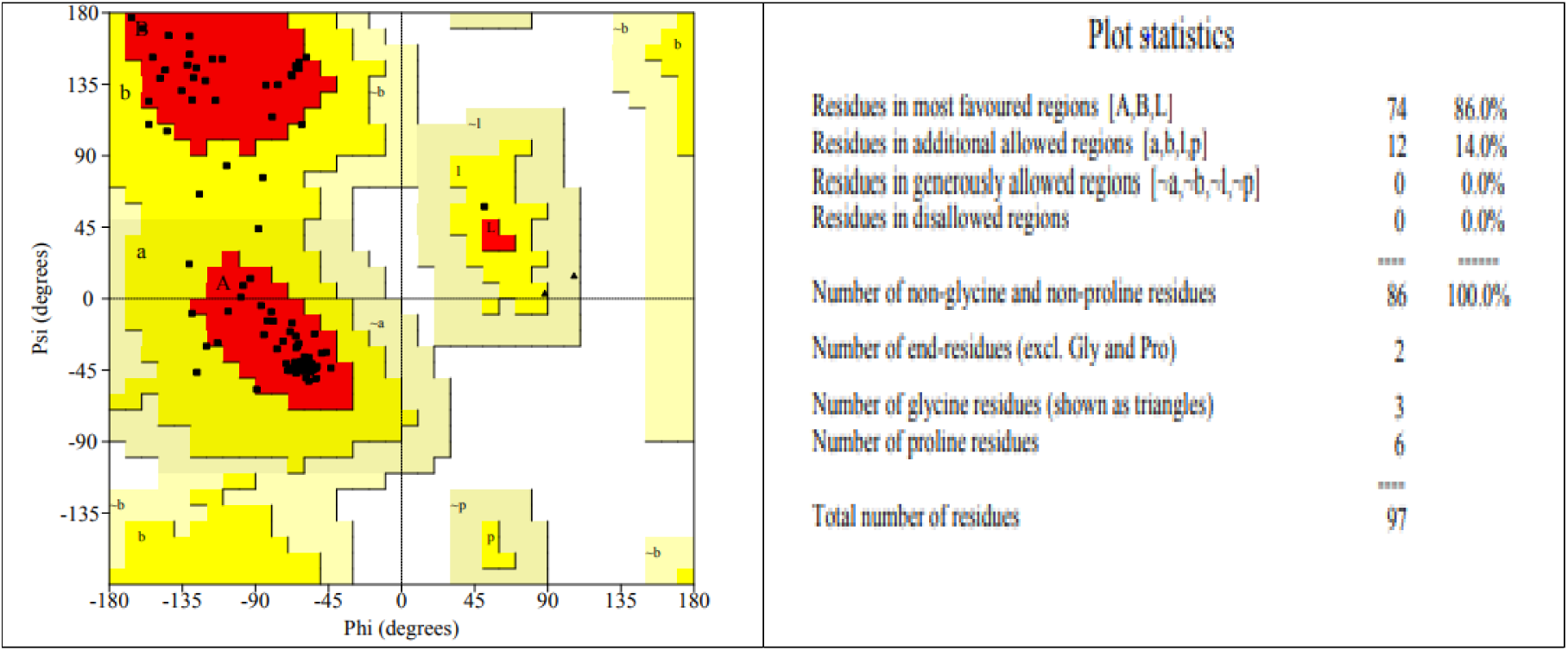
Ramachandran Plot using procheck server

**Figure6:**
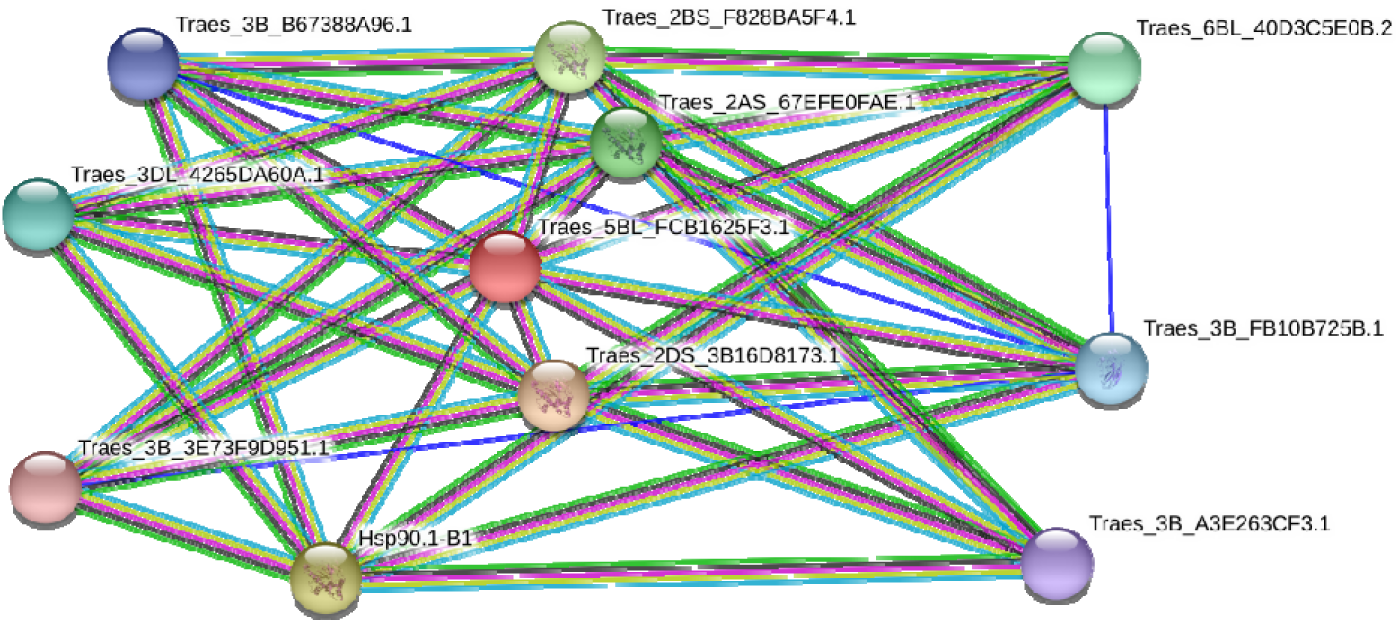
Protein-Protein Expression Using String Database.

## CONCLUSION

Thermal stress mainly heat stress may be a critical natural figure that antagonistically influencing trim surrender in numerous nations counting India. Plants are competent to adjust towards the high-temperature situations by transcriptional reconstructing, amid warm stretch. It was watched that heat stress moreover quickly actuates a set of heat stress protection genes, such as those encoding heat shock proteins. Heat shock factors (Hsfs) play a central administrative part in thermo resilience, here we recognized the potential heat shock factor protein from wheat, which can be utilized to create the transgenic plant with higher resilience towards thermal stress and eventually leads to a satisfactory nourishment supply. Confined Hsf2 is one of the major heat responsive translation variables that play a pivotal part in giving resistance against heat stress and controls the enactment of numerous metabolic pathways by anticipating the denaturation of numerous vital local proteins.

